# RNA is a critical element for the sizing and the composition of phase-separated RNA-protein condensates

**DOI:** 10.1101/457986

**Authors:** Marina Garcia-Jove Navarro, Shunnichi Kashida, Racha Chouaib, Sylvie Souquere, Gerard Pierron, Dominique Weil, Zoher Gueroui

## Abstract

Liquid-liquid phase separation is thought to be a key organizing principle in eukaryotic cells to generate highly concentrated dynamic assemblies, such as the RNP granules. Numerous *in vitro* approaches have validated this model, yet a missing aspect is to take into consideration the complex molecular mixture and promiscuous interactions found *in vivo*. Here we report the versatile scaffold “ArtiG” to generate concentration-dependent RNA-protein condensates within living cells, as a bottom-up approach to study the impact of co-segregated endogenous components on phase separation. We demonstrate that intracellular RNA seeds the nucleation of the condensates, as it provides molecular cues to locally coordinate the formation of endogenous high order RNP assemblies. Interestingly, the co-segregation of intracellular components ultimately impacts the size of the phase-separated condensates. Thus, RNA arises as an architectural element that can influence the composition and the morphological outcome of the condensate phases in an intracellular context.

## Introduction

Membrane-less organelles, by localizing and regulating complex biochemical reactions, are ubiquitous functional subunits of intracellular organization. Among them, ribonucleoprotein (RNP) granules, which include processing bodies (P-bodies), stress granules (SGs), germ granules, nucleoli, Cajal bodies, etc., are supramolecular assemblies of RNA and protein found in eukaryotic cells ^1^. They regulate RNA processing and thereby play a pivotal role in overall gene expression output, while their dysfunction is linked to viral infection, cancer and neurodegenerative diseases ^1,2^. Although RNP granules exhibit different compositions and functions depending on the cellular context, they have strikingly common features concerning their biophysical behavior and assembly process. RNP granules behave as highly concentrated liquid-like condensates that rapidly exchange components with the surrounding medium ^3,4^ and their formation relies on a self-assembly process ^5,6^. A general model has emerged where RNP granules generate from liquid–liquid phase separation, driven by low affinity interactions of multivalent proteins and/or proteins containing intrinsically disordered regions (IDRs) ^7–11^. However, new roles of RNA in the regulation of granule formation, which has been long assigned to protein components, are being uncovered ^12^. RNAs can promote phase separation synergistically with protein-protein interactions but also independently ^13,14^. *In vitro* studies identified that RNA modulates the biophysical properties of liquid droplets, by tuning their viscosity and their dynamics ^15,16^. By acting as a molecular seed, RNA contributes to the spatiotemporal regulation of phase-separated granules ^17^. For instance, the polar positioning of germ granules found in *C. elegans* reflects an RNA competition mechanism that regulates local phase separation ^18,19^, while rRNA transcription allows the cells to overcome the inherent stochastic nature of phase separation by timely seeding of nucleolus assembly in *D. melanogaster* embryos ^20,21^. More recently, it has been proposed that defects in nuclear RNA levels lead to excessive phase separation of IDR-containing RNA binding proteins (RBPs) such as FUS and TDP-43 ^22^. Moreover, RNA appears to template the molecular composition of the granules as shown for polyQ-dependent RNA–protein assemblies ^23^. In this context, the development of generic methods, integrating the knowledge accumulated from phase separation *in vitro* studies, would be particularly acute to elucidate the general principles of the structuring role of RNA within a condensed phase in the cellular environment.

In the present study, we describe the ArtiGranule (ArtiG) bottom up approach to form, within living cells, RNA-protein assemblies that recapitulate the features of phase-separated liquid condensates. First, through genetic engineering, we modified the oligomeric ferritin protein to generate a modular and versatile scaffold capable of self-interacting with low affinity. Upon reaching a critical concentration, this scaffold assembles into micrometer-sized ArtiG condensates within the cytoplasm of the cells. Then we built up the design with a canonical RNA binding domain to enable the biochemically neutral condensates to recruit endogenous RNAs. We implemented the PUM.HD domain of human Pumilio 1, a translational repressor that accumulates in P-bodies ^24^, and we demonstrated that the resulting ArtiG^PUM^ specifically recruit endogenous Pumilio 1 RNA-targets. Finally, this method enabled us to uncover the impact of intracellular RNA in different aspects of the condensate assembly: (i) ArtiG^PUM^ form more efficiently than control ArtiG, underlining that the recruitment of endogenous RNA seeds and facilitates the condensate nucleation. (ii) The size and polydispersity of ArtiG^PUM^ per cell is strikingly reduced, while their number is higher, compared to control ArtiG. This indicates that the incorporation of endogenous RNAs modulates the morphological outcome of phase-separated condensates. (iii) Micrometric bodies composed of P-body components localize at the periphery of ArtiG^PUM^, revealing that ArtiG^PUM^ subsequently and specifically co-segregate the RBPs associated to the Pumilio-targeted RNAs. Interestingly, the ArtiG^PUM^ interact with SG elements exclusively in response to stress. We suggest that the multivalent RNAs displayed on ArtiG^PUM^ surface act as molecular cues that seed the recruitment of specific subsets of RBPs/RNAs and coordinate the coexistence of endogenous higher-order assemblies, such as P-Body-like and SG-like assemblies. Furthermore, the docking of biochemically different phases, which is a conserved feature of numerous RNP granules, emerges as a parameter that can regulate the size of the condensates by limiting the growth by structural component addition or coalescence.

## Results

### ArtiG assemble into micrometric condensates within the cytoplasm of living cells by a concentration-dependent phase transition

The formation of RNA-protein condensates is thought to be driven by liquid-liquid phase separation through weak multivalent interactions between biomolecules ^8^. According to this model, the multivalent interactions promote the demixing, while their low affinity enables the components to dynamically exchange within the condensed phase and with the surrounding medium. Inspired by these findings, we took advantage of ferritin self-assembly in a 24-mer nanocage to generate a multivalent 3D scaffold, to which we added an interaction module. We engineered the light chain of human ferritin by fusing the F36M-FKBP (Fm) protein to the N-terminus of the ferritin monomer. The Fm protein is a point mutant of human FKBP protein that has the property of forming homodimers with micromolar affinity ^25^. The resulting chimeric Fm-ferritin (FFm) has 24 Fm proteins pointing outside the nanocage, which act as self-interacting domains. Thus, every FFm could undergo up to 24 interactions with other FFm nanocages with micromolar affinity. Such a protein scaffold can easily be functionalized by fusing a protein of interest (POI-FFm), for instance a fluorescent protein or a RNA binding domain (Figure 1A).

**Figure 1:**
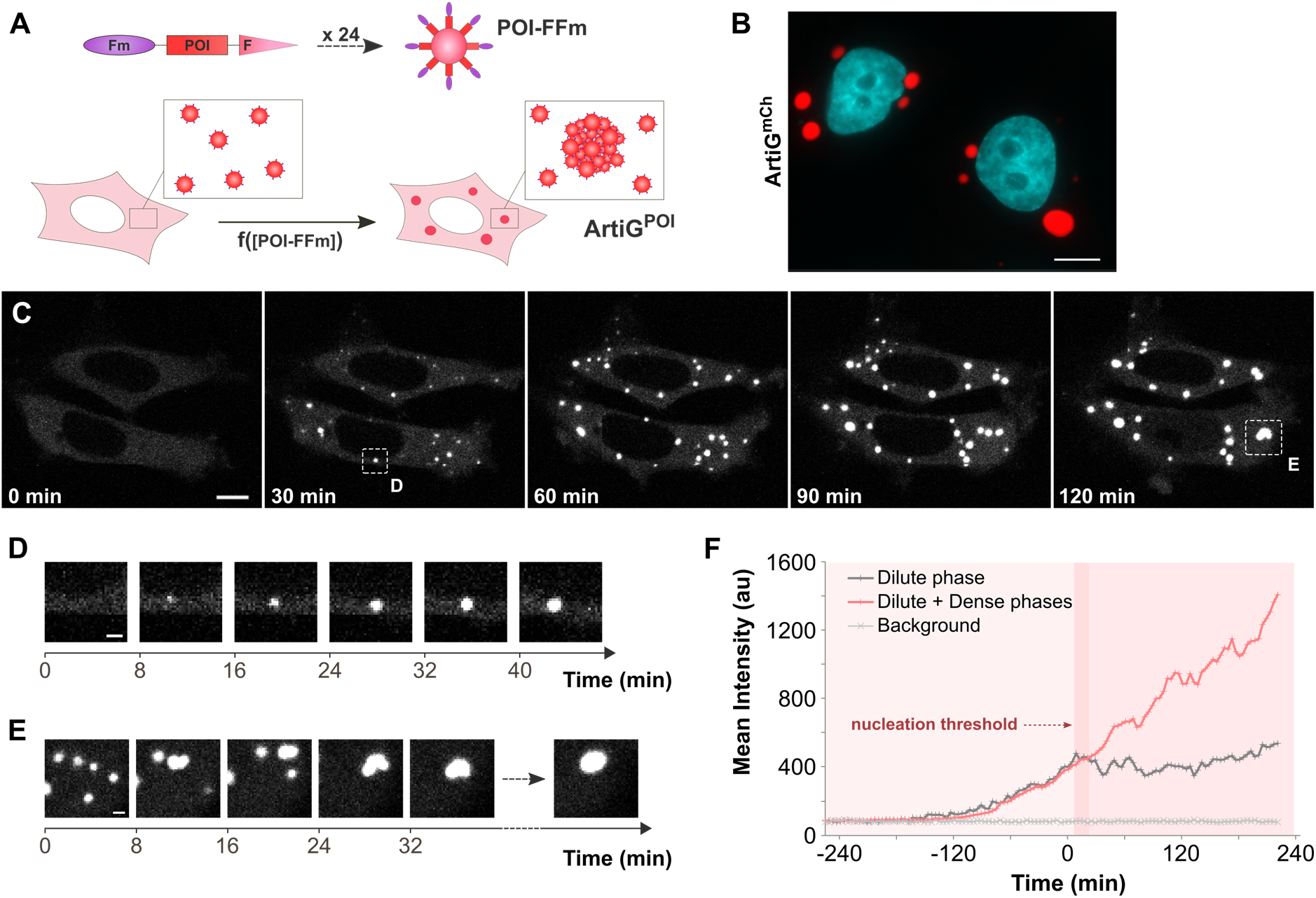
FFm scaffold forms concentration-dependent condensates in living cells. (**A**) Schematic of POI-FFm multivalent self-interacting scaffold. Genetically engineered ferritins assemble into ArtiG^POI^ through a phase separation process (Fm = F36M-FKBP, POI = Protein of Interest, F = Ferritin, ArtiG = ArtiGranule). (**B**) Representative confocal image of ArtiG^mCh^ (red) in HeLa cells, 24 h after transfection of mCherry-FFm construct. Nuclei were stained with Hoechst (blue). Scale bar, 10 μm. (**C**) Representative time-lapse confocal images of ArtiG^mCh^ (white) nucleation and growth in HeLa cells expressing mCherry-FFm construct. Scale bar, 10 μm. (**D**) Representative time-lapse confocal images of the growth of an isolated condensate. Scale bar, 2 μm. (**E**) Representative time-lapse confocal images of several ArtiG^mCh^ coalescing into a larger condensate. Scale bar, 2 μm. (**F**) Comparison of the temporal evolution of the dilute cytosolic mCherry-FFm fluorescence (dilute phase), with the total cytoplasmic fluorescence (dilute and dense phases) for the lower cell shown in (C). For the purpose of representation, the images in (C), (D) and (E) are slightly saturated.

To test the strategy, we first expressed an mCherry-FFm construct in HeLa cells. 24 hours after of transfection, about 30% of the transfected cells displayed a few micrometric fluorescent condensates dispersed throughout their cytoplasm, while the other cells exhibited homogeneous diffuse cytosolic mCherry-FFm fluorescence (Figures 1B and S1A). Neither fluorescence nor mCherry-FFm condensates were detected in the nucleus, in agreement with the diameter of the folded mCherry-FFm nanocages, which exceeds the size limit to passively diffuse through nuclear pore complexes. When Fm protein was replaced by the wild-type FKBP protein, which is unable to homodimerize, the resulting FKBP-mCherry-ferritin construct did not promote condensate formation and remained diluted in the cytoplasm of cells (Figure S1B). The formation of FFm condensates was recapitulated in HEK293 cells with equivalent results (Figure S1C). From now on, the mCherry-FFm condensates will be referred as ArtiG^mCh^ and more generally, any POI-FFm condensate as ArtiG^POI^ (Figure 1A).

In order to characterize ArtiG^mCh^ nucleation and growth steps, we next followed mCherry-FFm expression after 8-10 hours of transfection using time-lapse live imaging. At low expression levels, mCherry-FFm was diffuse in the cytoplasm of HeLa cells (Figure 1C, first panel). As mCherry-FFm concentration increased as a function of time, the nucleation of several micrometric fluorescent ArtiG^mCh^ condensates was observed (Figure 1C). The spherical ArtiG^mCh^ were homogeneously distributed and mobile within the cytoplasm (Movie S1). Individual condensates first exhibited a rapid growth phase both in size and fluorescence intensity, before slowing down after about 40-60 minutes (Figures 1D and S1D); while condensates that encountered one another, coalesced to form larger spherical bodies (Figure 1E and Movie S2). We monitored the accumulation of the dilute cytosolic mCherry-FFm fluorescence (dilute phase) and of the total cytoplasmic fluorescence (dilute + condensed phases) as a function of time (Figures 1F and S1E). Concurrently with the formation of the first ArtiG^mCh^, the cytosolic fluorescence intensity (dilute phase) stopped increasing to eventually reach a stationary phase, whereas the total cytoplasmic fluorescence intensity (dilute + condensed phases) kept increasing over time. These results suggest that the nucleation of ArtiG^mCh^ occurs at a critical concentration of cytosolic mCherry-FFm, a signature of first-order phase transitions. Furthermore, ArtiG^mCh^ seem to establish a dynamic equilibrium buffering the variations of FFm concentration in the cytoplasm. Taken together, these data illustrate that FFm scaffold undergoes a phase transition to form ArtiG micrometric condensates within the cytoplasm of living cells, in a concentration-dependent manner.

### ArtiG recapitulate the main characteristics of phase separated liquid droplets

A number of studies has highlighted that membrane-less organelles have specific material properties and behave as liquid or gel compartments ^26^. Therefore, we further assessed the morphological and biophysical characteristics of the ArtiG.

The ultrastructural characterization of ArtiG^mCh^, by electron microscopy of glutaraldehyde-fixed and Epon-embedded thin sections of the mCherry-FFm expressing cells, revealed micrometric round bodies not seen in control cells (Figure 2A). They were not delineated by a membrane and they were usually distant from cytoplasmic organelles such as mitochondria or the endoplasmic reticulum. In agreement with these observations, no fluorescent lipophilic staining was detected around ArtiG^EGFP^ in fluorescence microscopy (Figure S2A) and the different fluorescent ArtiG exhibited generally spherical shapes (Figures 1 and 2). Besides, ArtiG that deformed under mechanical stress relaxed back in a spherical shape (e.g. after being trapped between the nuclear and plasma membranes – Figure S2B). These dynamics and the spherical morphology indicate a liquid-liquid demixing state of the ArtiG.

**Figure 2:**
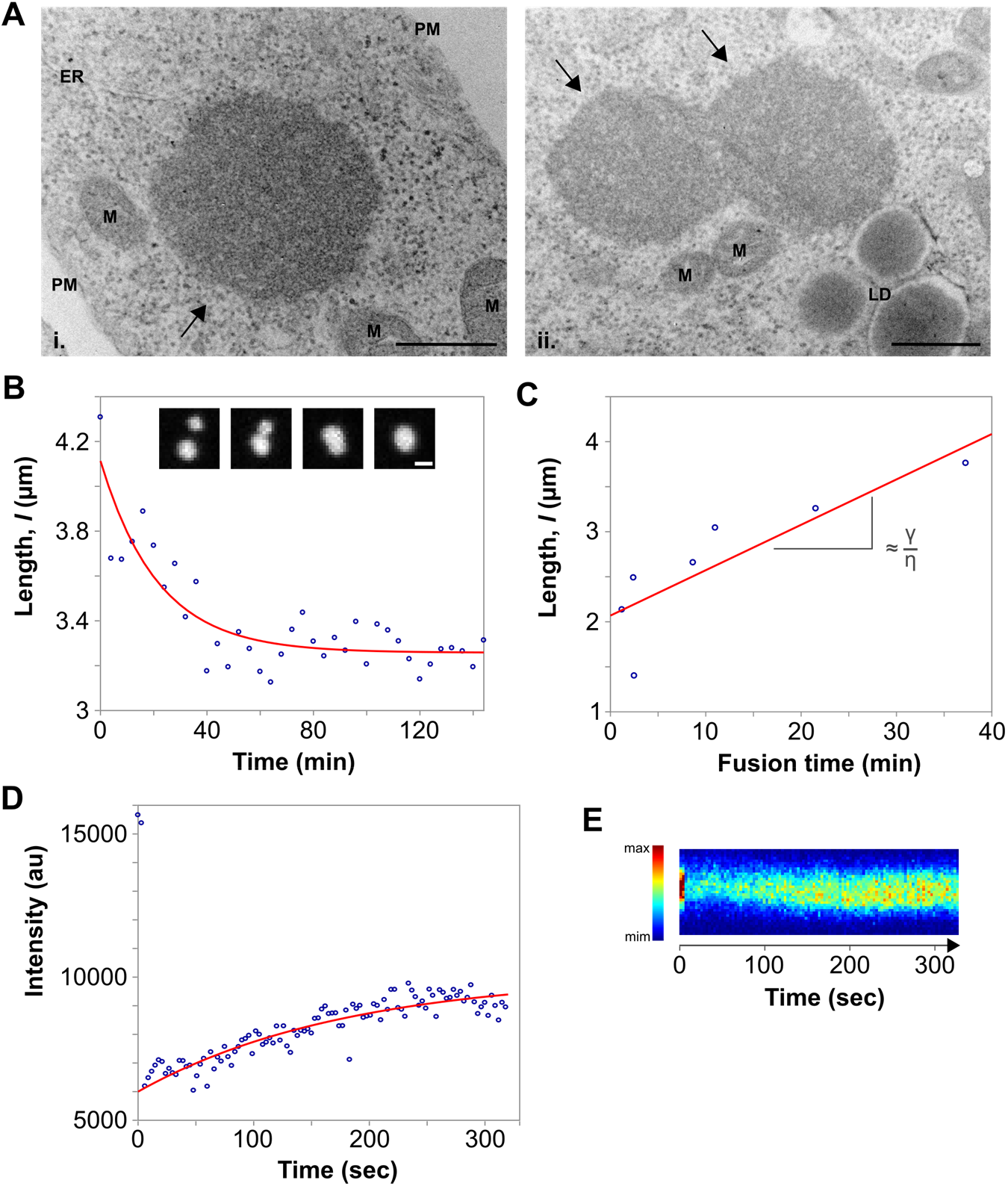
ArtiG^mCh^ recapitulate the main characteristics of phase-separated liquid droplets. (**A**) Ultrastructure of an individual ArtiG^mCh^ (i., black arrow) and of two ArtiG^mCh^ (ii., black arrows) undergoing fusion in HeLa cells, 24 h after transfection of mCherry-FFm construct. PM = Plasma membrane, ER = Endoplasmic reticulum, M = Mitochondrion, LD = Liquid droplet. Scale bar, 500 nm. (**B**) Temporal evolution of the distance between two ArtiG^mCh^ fusing with each other and relaxing into a single droplet within a characteristic time of 40 min. Insert: time-lapse of the corresponding fusion event. Scale bar, 2 μm. (**C**) Plots of the characteristic relaxation time of different fusion events as a function of the diameter, with the slope giving the capillary velocity (ratio between the surface tension γ and the viscosity ζ). (**D**) Example of recovery of fluorescence intensity after photobleaching of a 2 μm ArtiG^mCh^. (**E**) Kymograph representation of the fluorescence recovery of the ArtiG^mCh^ analyzed in (D).

An additional hallmark of phase separation displayed by ArtiG is their tendency to fuse. As observed in Figures 1D and 2B, nearby ArtiG^mCh^ tend to coalescence to form larger bodies, which could give rise to the cells displaying a few large spherical condensates 24 hours after transfection (Figures 1B and S1A). By computing the fusion time as a function of the diameter of the resulting ArtiG^mCh^ (Figure 2C) and by assuming that surface tension drives the fusion process whereas viscosity tends to impede it, we determined that ArtiG^mCh^ have an effective viscosity of about 10^3^ Pa.s (Methods). This is in the same order of magnitude than the effective viscosity measured for nucleoli in *Xenopus laevis* and for large RNP granules in *Caenorhabditis elegans* oocytes ^4,27^.

Finally, we performed fluorescence recovery after photobleaching experiments (FRAP) and found that, after photobleaching of the ArtiG^mCh^ components, a fraction of the fluorescent signal (35%) recovered with a timescale of about 2 minutes for micrometric granules (Figures 2D and 2E). This illustrates that ArtiG^mCh^ components are mobile and exchange with the surrounding cytoplasm.

Taken together, these data demonstrate that ArtiG^mCh^ recapitulate the hallmarks of liquid-like behavior described for native RNP granules, as they are spherical and relax into one spherical droplet after fusion and deformation, but also as ArtiG^mCh^ components dynamically exchange with the dilute cytosolic phase.

### ArtiG can be functionalized with a RNA-binding domain to recruit specific endogenous RNAs

To shed light on the impact of RNA on phase separation of RNA-protein condensates in a native cellular environment, we implemented a canonical RNA binding domain in our multivalent FFm scaffold. We chose the Pumilio homology domain (PUM.HD) of human Pumilio-1 protein as: (i) PUM.HD is a well-characterized RNA-binding domain ^28^; (ii) Pumilio-1 is known to bind specific RNA elements that accumulate in P-bodies ^24^, but do not promote their assembly when overexpressed in cells ^29^; (iii) By its RNA-binding activity, Pumilio-1 is a key regulator of numerous cellular processes, including translation repression ^30^. To generate PUM.HD-FFm self-interacting multivalent scaffold, PUM.HD was inserted between the Fm protein and the ferritin monomer (Figure 1A).

We first confirmed by co-expressing EGFP-FFm and mCherry-FFm constructs that cells could display ArtiG^EGFP/mCh^ hybrid condensates containing two different ferritins (Figure S3A). We next directed the formation of hybrid ArtiG^mCh^/^PUM^ by co-expressing mCherry-FFm and PUM.HD-FFm, in a 5:1 plasmid ratio, and we monitored the protein expression by western blot analysis (Figure S3B). We assessed the formation of ArtiG^mCh/PUM^ by time-lapse confocal microscopy 8-10 hours after transfection (Figure 3A). As for ArtiG^mCh^, the fluorescence was first diffuse in the cytoplasm before concentrating in several bright bodies that grew as a function of time in a concentration-dependent manner (Figures 3A and S3C). 24 hours after transfection, cells displayed several micrometric condensates (Figure 3B). On the other hand, when we transfected a FKBP-mCherry-PUM.HD-ferritin construct, which is multimeric but cannot self-interact, no fluorescent condensates were observed (Figure S3D). In order to validate the presence of RNA in ArtiG^mCh/PUM^, we analyzed the localization of a EGFP-fused Poly(A)-binding protein (PABP-EGFP) as a fluorescent reporter of polyadenylated RNAs. As shown in Figure 3C, when over-expressed, PABP-EGFP accumulated around ArtiG^mCh/PUM^, which exhibited an enriched corona of EGFP fluorescent signal, but not around control ArtiG^mCh^. Collectively, the co-expression of a RNA-binding PUM.HD-FFm scaffold with a neutral mCherry-FFm scaffold eventually leads to the concentration-dependent phase separation of micrometric protein condensates, which recruit polyadenylated RNAs.

**Figure 3:**
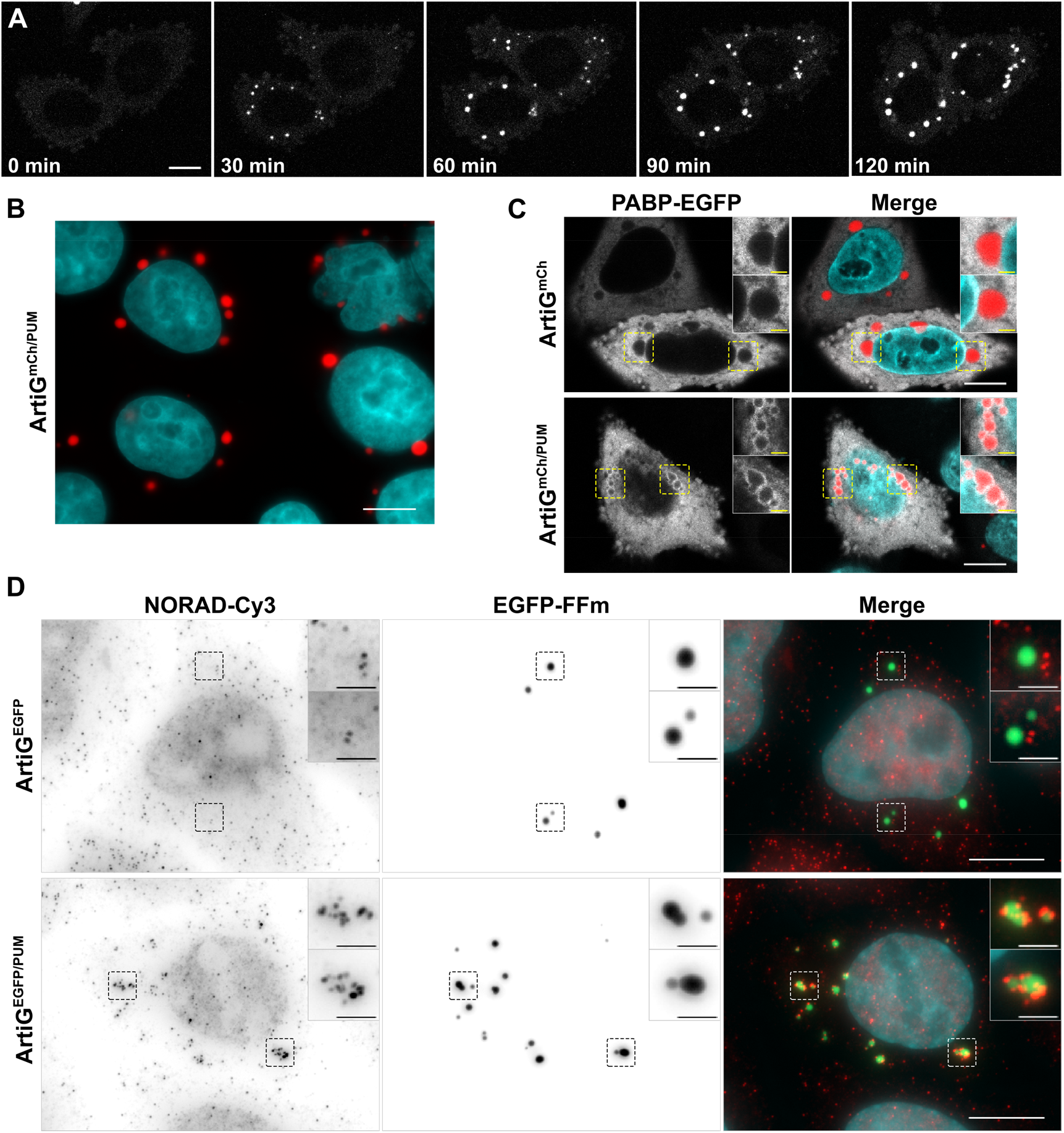
ArtiG^PUM^ recruit specific endogenous RNAs. (**A**) Representative time-lapse confocal images of HeLa cells expressing mCherry-FFm and PUM.HD-FFm constructs. Scale bar, 10 μm. For the purpose of representation, the images are slightly saturated. (**B**) Representative confocal image of ArtiGmCh/PUM (red) in HeLa cells, 24 h after transfection of mCherry-FFm and PUM.HD-FFm constructs. Nuclei were stained with Hoechst (blue). Scale bar, 10 μm. (**C**) To visualize polyadenylated RNA, a PABP-EGFP fusion was co-transfected with mCherry-FFm and PUM.HD-FFm (red) into HeLa cells 24 h before fixation. Confocal imaging. Scale bar, 10 μm. Zoom, 2 μm. (**D**) Epifluorescence imaging of anti-NORAD lncRNA smFISH-Cy3 (red in merge) in HeLa cells expressing ArtiG^EGFP^ (green in merge, upper row) and ArtiG^EGFP/PUM^ (green in merge, lower row). Nuclei were stained with DAPI (blue). Scale bar, 10 μm. Zoom, 2 μm.

We then investigated the specificity of RNA recruitment into ArtiG^mCh/PUM^. We focused on NORAD, a highly conserved and abundant long noncoding RNA (lncRNA), that has been described as a multivalent binding platform of PUM proteins and, by there, a key regulator of their activity ^31^. We examined the intracellular localization of endogenous NORAD lncRNA using the recently developed smiFISH (single-molecule inexpensive fluorescence in situ hybridization) technique ^32^. We found that ArtiG^EGFP/PUM^ were enriched in NORAD lncRNA (Figure 3D). Strikingly, NORAD-cy3 FISH signal was localized into micrometer-sized discrete patches on ArtiG^EGFP/PUM^. By contrast, ArtiG^EGFP^ were devoid of any NORAD-cy3 signal. To assess if RNA recruitment on the ArtiG^EGFP/PUM^ was specific, we performed a smiFISH to detect RAB13 mRNA, which is not a Pumilio 1-target and is known to localize at tips of protrusions ^33^. As expected, no colocalization between ArtiG^EGFP/PUM^ and RAB13 mRNA was observed (Figure S3E). Thus, these results show that ArtiG scaffold can be used to specifically target endogenous RNAs within a phase-separated condensate in cells.

### ArtiG^PUM^ recruit P-body components, have a higher frequency of nucleation and display a modified size

In a native context, the endogenous RNAs interact with specific RBPs that regulate their biogenesis and their fate. A recent study, unraveling the human P-body proteome and transcriptome, has reported that both Pumilio proteins and their RNA targets are significantly enriched in P-bodies ^24^. With this in mind, we assessed the presence of DDX6 and EDC4, as P-body markers, and ATXN2L as SG marker, in ArtiG^mCh/PUM^ by immunostaining cells 24 hours after transfection followed by confocal microscopy. Consistent with the neutral design of the ArtiG^mCh^ condensates, ArtiG^mCh^ did not colocalize with neither DDX6- or EDC4-labeled P-bodies (Figures 4A and S4A), nor with ATXN2L, even after arsenite-induced SG formation (Figures 4B and S4B). In contrast, ArtiG^mCh/PUM^ clearly displayed a DDX6 patchy corona on their surface (Figure 4A) and a similar result was obtained after EDC4 immunostaining, which formed micrometric patches attached to ArtiG^mCh/PUM^ (Figure S4A). This patchy organization around ArtiG was also observed by electron microscopy imaging. The immunogold detection of endogenous DDX6 on thin sections of ArtiG^mCh/PUM^ expressing cells reported a recurrent ultrastructural organization in distinct DDX6-labelled assemblies at the boundary of the condensates (Figure 4C). Additionally, ATXN2L fluorescent immunostaining of ArtiG^mCh/PUM^ expressing cells revealed that endogenous ATXN2L was mainly diffuse in the cytoplasm of the non-stressed cells (Figures 4B and S4B). Nevertheless, when cells were exposed to arsenite to induce *bona fide* SGs, micrometric ATXN2L patches were localized to ArtiG^mCh/PUM^ surface (Figures 4B and S4B). Altogether these data demonstrate that ArtiG^PUM^ selectively co-segregate P-body components in non-stressed cells, while SG components dock to ArtiG^PUM^ exclusively after arsenite stress.

**Figure 4:**
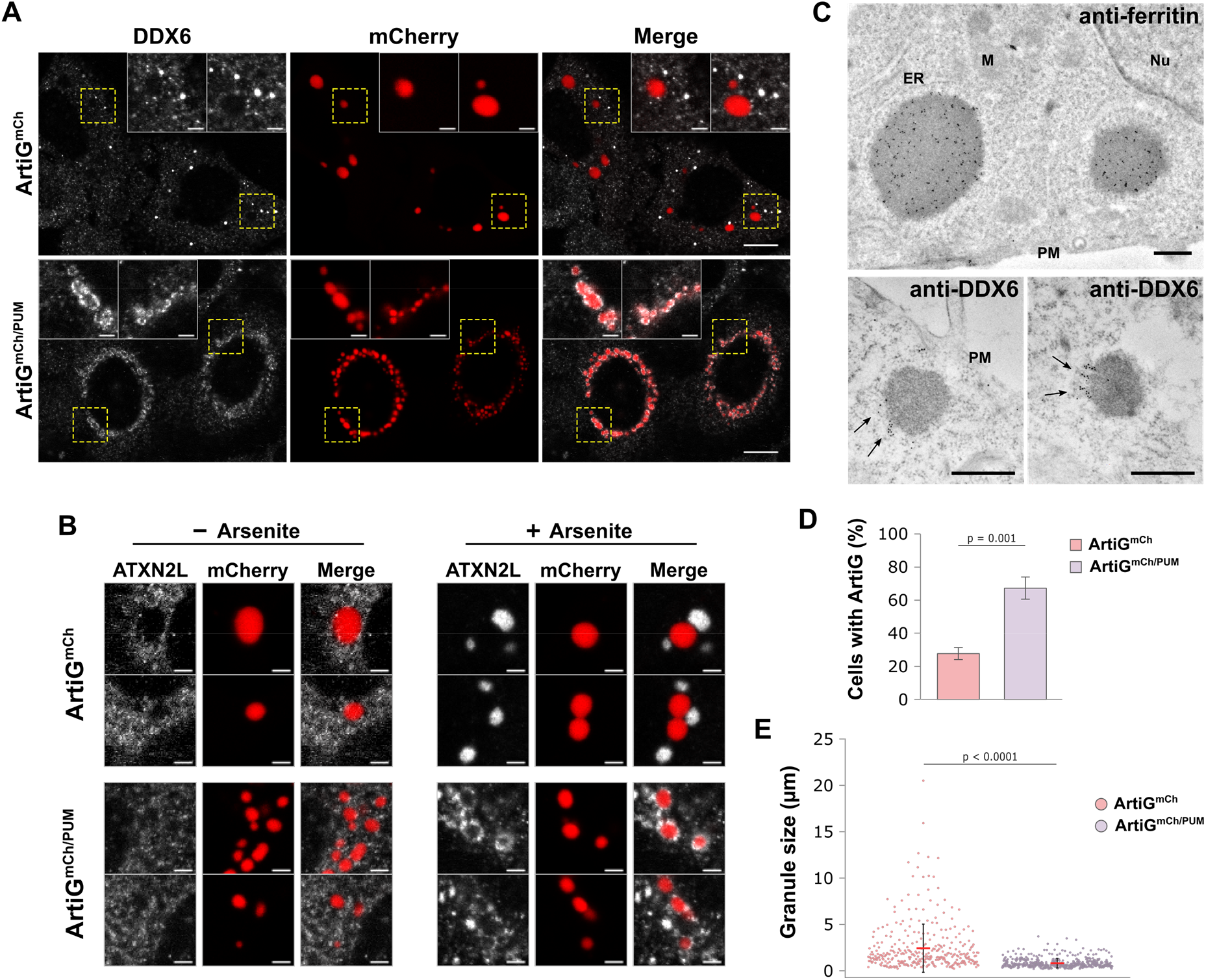
RNA modifies ArtiG^PUM^ composition, nucleation incidence and morphology. (**A**) HeLa cells expressing ArtiG^mCh^ and ArtiG^mCh/PUM^ (red) were fixed and analyzed by immunofluorescence using antibodies recognizing endogenous DDX6 (white). Confocal images. Scale bar, 10 μm. Zoom, 2μm. (**B**) HeLa cells expressing ArtiG^mCh^ and ArtiG^mCh/PUM^ (red) were treated or not with arsenite for 30 minutes, fixed and analyzed by immunofluorescence using antibodies recognizing endogenous ATXN2L (white). Enlarged regions of the confocal images presented in Figure S4B. Scale bar, 2μm. (**C**) As indicated, ferritin and endogenous DDX6 were identified in thin sections of ArtiG^mCh/PUM^ expressing cells, using a secondary antibody coupled to 10 nm gold particles. Ferritin is concentrated all-over ArtiG^mCh/PUM^ (upper panel), whereas DDX6 accumulates at the edge of the condensates (lower panels, black arrows). Scale bar, 500 nm. ER = Endoplasmic reticulum, M = Mitochondrion, Nu = Nucleus, PM = Plasma membrane. (**D**) Histogram of the percentage of transfected cells displaying ArtiG condensates 24 h after transfection of mCherry-FFm and PUM.HD-FFm constructs. Data represent the mean ± SD of three independent experiments. (**E**) Quantification of the size distribution of ArtiG^mCh^ and ArtiG^mCh/PUM^. Dots represent single measurements of ArtiG size and are pooled from three independent experiments. Means and SDs are superimposed.

Since ArtiG^PUM^ directly and indirectly incorporated components of endogenous RNP granules, we hypothesized that these endogenous components may modify the behavior of the synthetic condensates. First, we investigated the nucleation incidence of ArtiG^mCh/PUM^ in comparison with ArtiG^mCh^ 24 hours after transfection. ArtiG^mCh/PUM^ could be observed in ~70% of transfected cells, whereas ArtiG^mCh^ could only be detected in ~30% of transfected cells (Figure 4D), despite exhibiting similar protein levels (Figure S3B). Remarkably, ArtiG^mCh/PUM^ displayed a smaller size (mean ± SD = 0.83 ± 0.52 μm) and narrower size distribution (CV = 62 %) than ArtiG^mCh^ (2.44 ± 2.60 μm, CV = 107 %) (Figure 4E). Taken together, these results indicate that ArtiG^mCh/PUM^ establish a crosstalk with endogenous RNP components that organize in patchy assemblies around the ArtiG condensed phase. Furthermore, these interactions have a direct impact on the formation of the condensates, in terms of nucleation incidence and size.

### RNA as key regulator of composition, nucleation and sizing of the condensed phase

In order to assess more precisely the contribution of RNA in modifying the morphology and behavior of the composite condensates, we tuned the relative ratio of PUM.HD within the ArtiG. We transfected different plasmid ratios: 1:1, 5:1, and 10:1 of mCherry-FFm and PUM.HD-FFm constructs and protein level variation was confirmed by western blot analysis (Figure S5A). Strikingly, fluorescence microscopy image analysis revealed that ArtiG^mCh/PUM^ size and number scaled with the transfection ratio (Figure 5A). The more the ArtiG were enriched in RNA binding elements (PUM.HD-FFm), the more abundant and the smaller they were in the cells (Figures 5A and 5B). Regarding conditions with a transfection ratio of 1:1, cells displayed a large number of diffraction-limited ArtiG^mCh/PUM^, ~60 condensates per cell in average (of which 90 % was smaller than 0.4 μm in diameter, the diffraction limit of our optical setup). Electron microscopy acquisitions allowed the quantification of ArtiG^mCh/PUM^ size and dispersion (0.35 ± 0.15 μm, CV = 42 %) (Figure 5B). Interestingly, these sub-micrometric ArtiG^mCh/PUM^ coexisted without coalescing into larger condensates, despite their high concentration in the cytoplasm (Figure 5A, panel i.). In contrast, cells transfected with the plasmid ratio of 10:1 had a reduced number of ArtiG^mCh/PUM^ (in average ~12 condensates per cell) with a size and a distribution of 1.44 ± 1.39 μm (CV = 97 %), comparably to neutral ArtiG^mCh^ (~4 ArtiG^mCh/PUM^ per cell in average, 2.44 ± 2.60 μm, CV = 107 %) (Figures 5A, panels iii. and iv., and 5B). In this condition, ArtiG^mCh/PUM^ fused to relax into larger condensates (Movie S3). Finally, as described before, the transfection ratio of 5:1 led to cells exhibiting an intermediate phenotype (~19 ArtiG^mCh/PUM^ per cell in average, with 0.83 ± 0.52 μm in size, CV = 62 %) (Figures 5A, panel ii., and 5B). Single molecule detection of NORAD lncRNA confirmed that, despite the drastic size changes, all the ArtiG^mCh/PUM^ (1:1, 5:1 and 10:1) retained the capacity to bind RNA (Figures 5C and S5B). The recruitment of RNA in ArtiG^mCh/PUM^ was correlated to the specific recruitment of P-body elements in patchy micro-assemblies at the surface of the condensates (Figure S5C). Even in the most extreme condition of transfection we tested (ratio of 10:1), we still observed that ArtiG^mCh/PUM^ recruited RNA and P-body elements, indicating that even in minute amounts, the presence of PUM.HD-FFm affected the composition and behavior of the granules. These results suggested that the “RNA binding capacity” and thereby the RNA could directly modulate the morphologic fate of the phase-separated condensates in regards of size and number.

**Figure 5:**
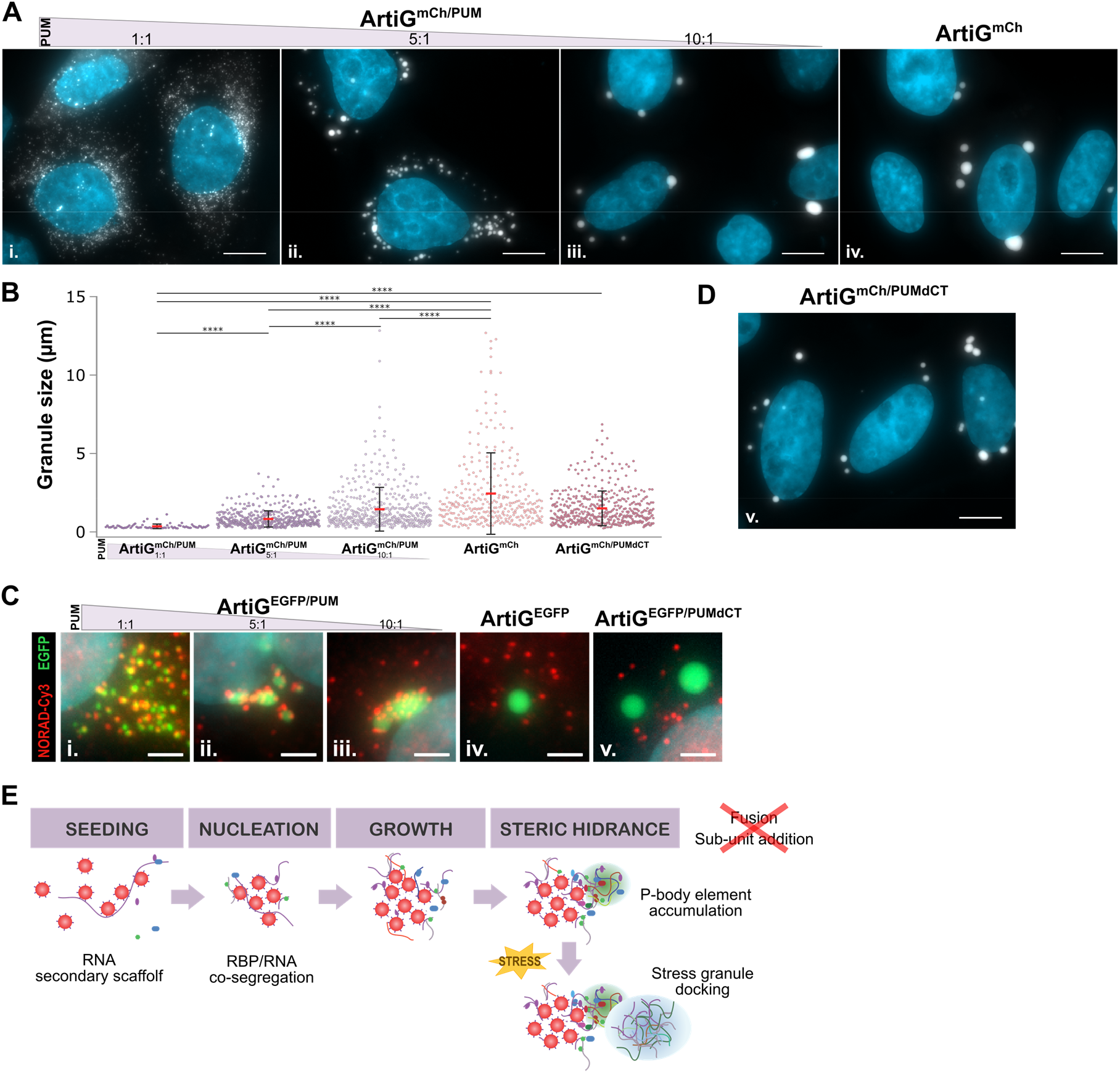
RNA is a critical element for ArtiG^PUM^ size scaling. (**A**) Representative confocal images of ArtiG^mCh/PUM^ (white) in HeLa cells, 24 h after transfection of mCherry-FFm and PUM.HD-FFm constructs in a plasmid ratio of 1:1 (i.), 5:1 (ii.), 10:1 (iii) and 1:0 (iv.). Nuclei were stained with Hoechst (blue). Scale bar, 10 μm. (**B**) Quantification of the size distribution of ArtiG^mCh/PUM^ and ArtiG^mCh^ described in (A). Dots represent single measurements for ArtiG size and are pooled from three independent experiments. Means and SDs are superimposed. **** p < 0.0001 For representation, the two largest ArtiG^mCh^ (> 15 μm, as shown in Figure 4E) were not included. (**C**) Epifluorescence images of anti-NORAD lncRNA smFISH-Cy3 (red) in HeLa cells expressing ArtiG^EGFP/PUM^, ArtiG^EGFP^ and ArtiG^EGFP/PUMdCT^ (green). The panels show enlarged regions of the cells represented in Figure S5B. Nuclei were stained with DAPI (blue). Scale bar, 2 μm. (**D**) Representative confocal image of ArtiG^mCh/PUMdCT^ (white) in HeLa cells, 24 h after transfection of mCherry-FFm and PUMdCT-FFm constructs in a plasmid ratio of 1:1. Nuclei were stained with Hoechst (blue). Scale bar, 10 μm. (**E**) Schematic of the model for ArtiG^PUM^ nucleation, growth and co-segregation of endogenous RNPs.

To further validate the role of RNA in shaping the condensate phase, we constructed a deletion mutant of PUM.HD-FFm in order to reduce its RNA binding capacity. PUM.HD is a sequence-specific RNA binding domain, which is characterized by eight tandem imperfect copies of a structural motif (R1 to R8), that respectively interact with a different RNA nucleotide of the targeted RNA-sequence ^28,34^. We hypothesized that by truncating the C-terminal region of PUM.HD (mutant PUMdCT), we could weaken its RNA-binding capacity and hence the sizing phenotype of ArtiG^PUM^. After confirming the assembly of ArtiG^mCh/PUMdCT^ mutant condensates in living cells (Figures 5D and S5D for western blot analysis), we next assessed that the deletion of the last two motifs, R7 and R8, was sufficient to impair the RNA-binding capacity of PUM.HD. Colocalization experiments reported that ArtiG^mCh/PUMdCT^ were not able to recruit neither NORAD lncRNA (Figures 5C, panel v., and S5B), nor cytoplasmic PABP-EGFP at their surface (Figure S5E). Moreover, in the same transfection conditions (1:1 plasmid ratio with mCherry-FFm, Figure S5D for western blot analysis), ArtiG^mCh/PUMdCT^ displayed larger size and size distribution than wild-type ArtiG^mCh/PUM^: 1.50 ± 1.11 μm (CV = 74 %) versus 0.35 ± 0.15 μm (CV = 42 %) (Figure 5B). Again, the number of condensates per cell was inversely correlated with size, since cells exhibited ~10 ArtiG^mCh/PUMdCT^ per cell in average. These results demonstrated that the recruitment of RNA, and the resulting recruitment of its interacting partners, is the critical element responsible for the morphological changes of the phase-separated ArtiG^PUM^.

Altogether our data show that by conferring the capacity to recruit intracellular RNA to an initially inert scaffold that can undergo phase separation, we strongly modify the outcome of the condensates in terms of size and number. This illustrates how, in the context of an intracellular phase-separation, RNA is a critical architectural component in addition to its function as a carrier of genetic information.

## Discussion

Many of the biophysical hallmarks of phase separation model for RNP assembly have emerged from *in vitro* reconstitution experiments ^7,8^. Nevertheless, the study of phase separation processes in regards to molecular crowding, promiscuity, post-translational modifications, etc., requires the development of novel tools operating within the native intracellular environment ^35–38^. Inspired by pioneer experiments on liquid-liquid phase separation of multi-domain macromolecules ^8^, we engineered a multivalent self-interacting protein-scaffold, with the goal of being biochemically inert regarding the cellular environment. Our approach differs from strategies using protein scaffolds enriched with IDPs, which are very interesting for understanding, for instance, the role, promiscuity and aging of IDP assemblies, but which adds a supplementary level of complexity when working in the native context of cells as they undergo unrestrained homotypic interactions.

Our results show that our ArtiG approach forms micrometric assemblies that recapitulate the features of phase-separated condensates (Figure 1). Once a critical concentration of protein scaffold is reached in the cytoplasm, ArtiG start nucleating. As the ArtiG assemble, a dynamic equilibrium is established in which the concentration of the surrounding cytosolic phase is maintained constant, at the critical concentration level, through a buffering process by the condensed phase. The condensed phase incorporates the amount of scaffold continuously supplied by the cell and consequently grows in size (Figures 1F, S1E and S3C). This buffering property of phase separated compartments has been discussed as a possible role for intracellular condensates in maintaining the robustness needed for biochemical reactions, despite biological fluctuations ^5^. These hallmarks make our system reliable for testing predictions and studying phase separation in an intracellular environment.

We used ArtiG approach to examine the contribution of RNA elements to the formation of a condensed phase within a living system. We generated a simplified archetypal RNA-binding protein-condensate by incorporating a unique RNA-binding domain: PUM.HD, from human Pumilio-1 protein. Our minimal bio-mimicking system succeeds in assembling concentration-dependent RNA-protein condensates within the cytoplasm of living cells and in specifically recruiting Pumilio-1 RNA-targets (Figure 3). Interestingly, the presence of RNA in ArtiG^mCh/PUM^ correlates with a higher incidence of nucleation for these RNA-protein condensates, compared to purely protein-based ones (ArtiG^mCh^) (Figure 4D). Recent studies have proposed that RNA can provide spatiotemporal information and acts as a multivalent template that favors granule seeding ^16,17,21,23,39^. By carrying multiple binding sites for PUM proteins, NORAD lncRNA functions as a multivalent binding platform that sequesters a significant fraction of the cytosolic PUM proteins ^31^. This type of architecture would confer the capacity to localize local high-concentration assemblies that trigger the nucleation of a condensed phase to RNA. Altogether, these observations support that RNA can act as key regulator of *in vivo* phase separation by seeding the process ^18,21,22,40^.

An open question is how the biochemical composition of RNP granules is determined and maintained ^23^. Our results show that micrometric bodies of specific compositions are localized at the periphery of ArtiG^PUM^. The addition of a RNA binding domain to the protein-scaffold confers the capacity to communicate with intracellular components to the initial biochemically inert ArtiG condensates. We could imagine that by exposing and concentrating RNA locally, the ArtiG^PUM^ may recruit RBPs in a non-specific manner or induce SG formation by RNA mislocalization. In contrast, the RNA recruited by the ArtiG^PUM^ specifically co-segregates P-body components, such as DDX6 and EDC4, generating peripheral patchy DDX6- and EDC4-enriched bodies (Figure 4A, 4C and S4A). Two non-exclusive models could be envisioned: either the interaction of P-body components with the RNA may occur before its recruitment into the ArtiG^PUM^, or through the recruitment of a particular subset of RNAs, ArtiG^PUM^ expose molecular cues that spatially specify the nucleation P-body components (Figure 5E). This last picture is coherent with recent studies suggesting that RNP granules form by combinatory intermolecular RNA-protein, RNA-RNA and protein-protein interactions ^12, 41–44^. It is interesting to note that, in our system, two biochemically different phases, ArtiG and P-body-like, which share certain components, can coexist apparently immiscibly, as it has been observed for Cajal bodies with attached B-snurposomes ^45^. The local coordination of immiscibly condensed phases is even more striking under stress conditions, where we observe SGs interacting with the ArtiG^PUM^ (Figures 4B and S4B). Similarly, a physical juxtaposition of SGs and P-bodies is also observed when cells are stressed ^46–50^. As is seen for the natural granules, under stress conditions ArtiG^PUM^ may share transient interactions with SGs, physically linking the assemblies and leading to the docking of the compartments (Figure 5E).

There is a mounting interest in examining the mechanisms underlying size control of organelles during cell growth and embryonic development ^51,52^. Indeed the size of self-organized structures, such as the meiotic spindle or the centrosomes, scales with the cell dimension ^53^. One hypothesis is that size control could be mediated by the limiting supply of structural components found within the cell volume ^54–57^. Interestingly, intracellular phase transitions provide additional physical bases with which to understand the dependence of membrane-less organelle assembly, number and size, as a function of the cellular volume ^58,59^. We have been able to compare an archetypal phase separation of inert multivalent scaffolds to the phase separation of multivalent scaffolds that recruit a subset of intracellular components. ArtiG^mCh^ nucleation exhibits a concentration-dependence typical of first order phase transitions: upon reaching a critical concentration, multiple high-concentration assemblies emerge from the cytosolic dilute phase and grow in size, buffering the supply of scaffold subunits (Figure 1). In a simplified picture, a multi-droplet polydisperse mixture will coarsen by Ostwald ripening to form a single droplet, as the steady-state system tends to favor a few large condensates coexisting with the dilute phase ^60^. Yet, our observations suggest that, in an intracellular context, the growth of the ArtiG^mCh^ by coalescence dominates over Ostwald ripening. Ultimately, the first-assembled micrometric condensates will grow and coalesce in 1 or 2 large ArtiG^mCh^. In contrast, a large number of size-limited ArtiG^mCh/PUM^ are capable of coexisting within the cell. The RNA recruitment by the ArtiG^mCh/PUM^ has clear consequences for the growth and the polydispersity of the condensates: ArtiG^mCh/PUM^ growth process stops at some point, as a function of the amount of co-segregated biomolecules (Figure 5). The growth-limited phenotype of ArtiG^PUM^ was rescued when wild type PUM.HD was replaced by a mutant with defective RNA-binding activity. RNA appears to be the physical scaffold responsible for this feature. Still, why do ArtiG^mCh/PUM^ exhibit a limited size with respect to the protein-based ArtiG^mCh^? Theoretical studies suggest that coarsening by Ostwald ripening could be counterbalanced if non-equilibrium chemical reactions are present in the droplets ^61–63^. In *X. laevis* oocytes, nuclear actin network has been shown to stabilize nucleoli against coalescence ^64^. Recent work in yeast provides a model to explain the size of P-bodies from the intrinsic interactions of P-body components ^65^. In our system, we hypothesize that steric hindrance at the surface of ArtG^mCh/PUM^ might impede growth and coalescence (Figure 5E). The recruitment of endogenous RNA and RBP complexes at the surface of ArtG^mCh/PUM^ may reduce the subsequent sub-unit addition, thus limiting individual ArtiG^mCh/PUM^ growth. In addition, this recruitment possibly introduces enough spacing between nearby ArtiG^mCh/PUM^ to obstruct fusion events. Along with this scheme, our results show that the proportion of extrinsic interactions, which the artificial condensates make with intracellular components, scales the size of the final assemblies.

This work illustrates the interest of developing bottom-up approaches to dissect the general principles of RNP granule formation and function within living cells, taking into account the intracellular complexity of composition and interactions. Our work suggests that RNA can act as molecular cue that not only spatially seeds higher-order RNP assemblies, but also specifies their ultimate composition. Furthermore, the docking of biochemically heterogeneous phases arises as a critical parameter that affects the size of the condensates by limiting the growth by component addition or coalescence.

## Acknowledgments

The authors acknowledge all the members of the “Pole de Chimie Biophysique”, A. Hubstenberger, D. O. Wang, H. Saito and J.F. Joanny for fruitful discussions. We particularly thank A. Blanch Jover, E. Ipenday and M. Bénard for their help along the project, as well as A.G. Tebo and A. Colin for carefully reading the manuscript. We also thank M. W. Hentze and T. Fujita for kindly providing PABP-EGFP and Pumulio 1 plasmids.

M.G.J.N was supported by FRM (ING20150532742). S.K was supported by a French government fellowship, R.C. by ANR (ANR-14-CE09-0013-01).

This work was supported by the CNRS, Ecole Normale Supérieure, and the HFSP Program Grant (RGP0050/2014) to Z.G. and ANR (ANR-14-CE09-0013-01) to D.W.

## Author Contributions

M.G.J.N and Z.G conceived and analyzed the experiments. M.G.J.N carried out and analyzed the experiments. S.K contributed to the initial design of the experiments. R.C contributed to the implementation of FISH experiments. S.S and G.P performed and analyzed TEM experiments. D.W contributed to implement FISH and IF experiments and analyzing experiments.

M.G.J.N and Z.G wrote the manuscript and all authors were involved in revising it critically for important intellectual content.

